# A modular platform for bioluminescent RNA tracking

**DOI:** 10.1101/2022.07.02.498144

**Authors:** Kevin K. Ng, Kyle H. Cole, Lila P. Halbers, Christelle E.T. Chan, Erin B. Fuller, Chelsea Callicoatte, Mariajose Metcalfe, Claire C. Chen, Ahfnan A. Barhoosh, Edison Reid-McLaughlin, Alexandra D. Kent, Oswald Steward, Andrej Lupták, Jennifer A. Prescher

## Abstract

A complete understanding of RNA biology requires methods for tracking transcripts *in vivo*. Common strategies rely on fluorogenic probes that are limited in sensitivity, dynamic range, and depth of interrogation, owing to their need for excitation light and tissue autofluorescence. To overcome these challenges, we developed a bioluminescent platform for serial imaging of RNAs. Small RNA tags were engineered to recruit light-emitting luciferase fragments (termed RNA lanterns) upon transcription. Robust photon production was observed for RNA targets both in cells and in live animals. Importantly, only a single copy of the tag was necessary for sensitive detection, in sharp contrast to fluorescent platforms requiring multiple repeats. Overall, this work provides a foundational platform for visualizing RNA dynamics from the micro to the macro scale.

## Introduction

RNA dynamics play pivotal roles in a multitude of cellular processes (*1*). While we have a deep, molecular-level understanding of many facets of RNA biology *in vitro*, the picture in physiologically authentic environments—live animals—remains incomplete. This is due, in part, to a lack of methods for noninvasive tracking of RNAs *in vivo*. Conventional approaches rely on RNA tags coupled with fluorescent probes (*2-4*). Such platforms require excitation light, which can be difficult to deliver in whole organisms without invasive procedures, excision of tissues, or delivery of fluorogenic dyes (*5-7*). Furthermore, external light can induce autofluorescence, precluding sensitive detection of low abundance targets. Short imaging times are also necessary to avoid light-induced damage. Consequently, tracing the lifecycle of key RNAs in real time, in live mammals, has been elusive.

We reasoned that a potentially more suitable platform for RNA imaging in live animals could be achieved with bioluminescence. This modality relies on photon production from luciferase enzymes and luciferin small molecules. Since no excitation light is required, bioluminescence can provide superior signal-to-noise ratios *in vivo* for visualization of low-copy transcripts. Additionally, serial imaging is possible without concern for phototoxicity or tissue damage. The development of luciferases with higher photon outputs and improved thermal stability (e.g., NanoLuc and related variants) has enabled facile visualization of cells, biomolecules, and other features both on the micro and macro level (*8-11*). Recently, a split variant of NanoLuc was applied to RNA targets, setting the stage for precise detection of cellular transcripts (*12*).

Here we report a general method that leverages advances in bioluminescence technology for multi-scale RNA detection. The approach features split fragments of NanoLuc (NanoBiT) fused to MS2 and PP7 bacteriophage coat proteins (MCP and PCP), fusions that we have termed RNA lanterns. MCP and PCP bind distinct RNA aptamers (MS2 and PP7, respectively) that can be appended to transcripts of interest (*13, 14*). Upon transcription, MCP and PCP bind the MS2-PP7 containing RNA bait, bringing the luciferase fragments into proximity to assemble a functional, light-emitting enzyme. We extensively optimized both the lanterns and RNA bait to maximize signal turn-on and minimize the size of the protein-RNA complex. Notably, a single rigidified RNA was sufficient for sensitive imaging both in cells and *in vivo*, rendering much larger RNA tags unnecessary. Additionally, the RNA bait is modular and can be used in conjunction with other split luciferases and for multi-scale imaging. The tools reported here are thus immediately useful to studies of RNA dynamics.

## Results

### General strategy for visualizing RNAs with bioluminescent light

The overall strategy was dependent on three key steps: RNA bait formation, MCP/PCP binding, and NanoBiT complementation. We took inspiration from MCP- and PCP-based reporters comprising split fluorescent proteins to visualize transcripts (*15*). We envisioned that the RNA-binding proteins could be fused to the NanoBiT system (*16*), which has been used extensively to examine protein-protein interactions and other biomolecular networks (*17-19*). We appended each part of NanoBiT (the short peptide—SmBiT—and the larger protein fragment—LgBiT, **Fig. 1A**) to MCP and PCP, respectively. The resulting fusions (termed RNA lanterns) were hypothesized to bind their cognate aptamers on a single transcript, inducing NanoBiT complementation and generating photons in the presence of luciferin. The requisite components had not been used together prior to this work, necessitating optimization of each step.

**Fig. 1.**
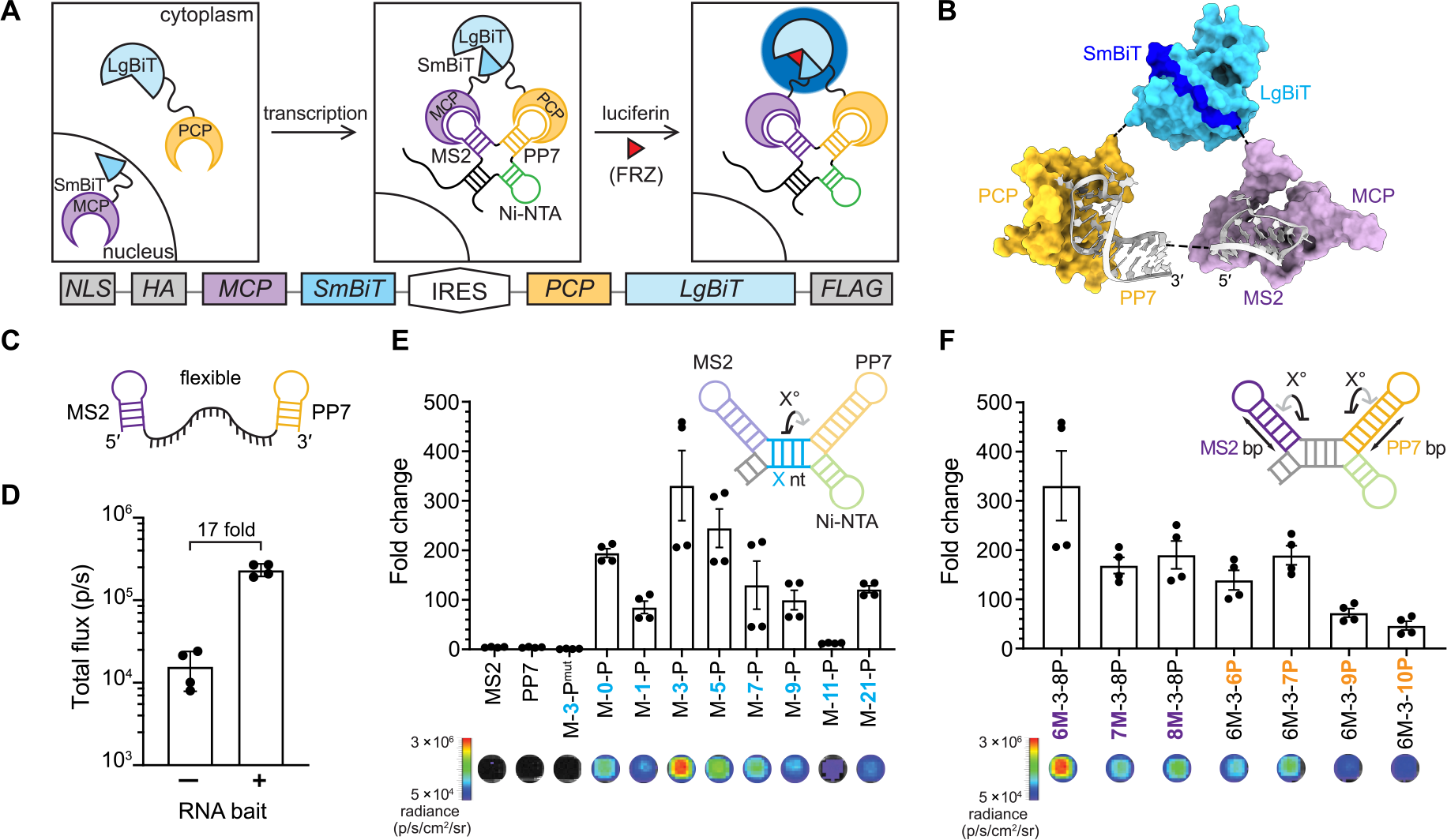
An optimized platform for tracking RNA dynamics. (**A**) Overall strategy to visualize transcripts using RNA lanterns. The lanterns comprise fusions of MS2 coat protein (MCP) and PP7 coat protein (PCP) with NanoBiT fragments (SmBiT and LgBiT, respectively). Transcription of bait RNA (comprising MS2 and PP7 aptamers) drives NanoBiT heterodimerization. In the presence of a luciferin substrate (furimazine, FRZ), light is produced (blue glow). The bicistronic construct encoding the RNA lantern is shown below the scheme. MCP and PCP were fused with HA and FLAG tags, respectively, for expression analyses. (**B**) Modeling of the RNA lantern complex. Crystal structures of MS2 (purple, 1ZDI) (*32*), PP7 (orange, 2QUX) (*33*) and NanoLuc (5IBO), highlighting LgBiT (dark blue) and SmBiT (cyan), were modeled in ChimeraX (*20*). The MCP-SmBiT/PCP-LgBiT complex was modeled by aligning the corresponding N- and C-termini of each protein and aligning the 5′ and 3′ ends of the aptamers. (**C**) Predicted secondary structure of flexible RNA bait, as calculated by RNAfold (*21*). (**D**) Initial tests of RNA lantern with a flexible RNA bait via an *in vitro* transcription and translation (IVTT) assay. The total flux observed in the absence or presence of RNA bait is plotted. (**E**) Engineered rigid RNA baits provided robust photon output in IVTT assay. RNA baits containing varying spacers between the MS2 and PP7 aptamers were constructed, with spacer length (X nt, blue, denoted below each bar graph), modulating the distance and helical phase angle between the aptamers. Fold change in signal over RNA lantern alone is plotted. MS2, PP7, and M-3-P^mut^ denote baits comprising isolated aptamers or a mutated PP7 aptamer (fig. S3, A and B), respectively, none of which were expected to result in RNA lantern assembly. Representative luminescence images are shown below the graph. (**F**) Rigid RNA baits comprising various MS2 and PP7 stem lengths (top right, purple and orange for the MS2 and PP7 aptamers, respectively). Luminescence readouts were acquired following IVTT. Fold change in signal over RNA lantern alone is plotted. Representative luminescence images are shown below the graph. Error bars represent the standard error of the mean (SEM) for *n* = 2 replicates (D) and *n* = 4 replicates (C, E, and F).

As a starting point, we designed a bicistronic construct with an internal ribosome entry site (IRES) to mediate co-expression of the RNA lantern components (**Fig. 1A**). MCP was fused to SmBiT and PCP was fused to the larger protein fragment (LgBiT), building on a previously published split fluorescent protein platform, in which MCP and PCP were fused to the N- and C-terminal fragments of the Venus fluorescent protein, respectively (*15*). MCP was further tagged with a nuclear localization signal (NLS) to reduce background complementation. This same strategy was used previously for the split Venus system (*15*) to separate the lantern fragments in the absence of the RNA bait. Upon expression, transcripts fused to the RNA bait could transport an MCP-fusion from the nucleus into the cytoplasm (or capture *de novo* translated MCP in the cytoplasm). Eventual binding of the PCP fusion would co-localize both halves of the RNA lantern on the target transcript, enabling NanoBiT complementation and thus light production. Additionally, the NLS reduces the likelihood of non-specific SmBiT/LgBiT complementation, a problem encountered in previous studies that resulted in diminished sensitivity (*12*).

We modeled the RNA lanterns using ChimeraX (*20*) to assess the design of the fusions (**Fig. 1B**). No obvious steric clashes or unfavorable orientations with a 5′-MS2-PP7-3′ RNA bait were observed, suggesting that lantern assembly was possible. NanoBiT complementation appeared feasible even with juxtaposed MS2 and PP7 aptamers. Small, compact baits are attractive tags to avoid disrupting the structures or functions of target RNAs. Encouraged by the modeling results, we moved forward with the lantern design and anticipated that additional engineering of the lantern components (linkers and orientation) would be necessary to maximize luciferase complementation and signal output.

### Biochemical optimization of the RNA tag and lanterns

The RNA detection platform relies on efficient NanoBiT formation, which can be tuned based on the SmBiT peptide sequence (*16*). We examined two SmBiT peptides (SmBiT^high^, *K*_D_ = 180 nM; SmBiT^low^, *K*_D_ = 190 µM) to determine which would provide the best dynamic range: minimal signal in the absence of RNA bait and robust signal in its presence (**fig. S1A**). The designer probes were first expressed using an *in vitro* transcription/translation system (IVTT) featuring transcription by T7 RNA polymerase and translation in rabbit reticulocyte lysate (RRL). Background signal was determined in the absence of RNA bait. Full complementation of translated NanoBiT was achieved by adding exogenous SmBiT^ultra^ (*K*_D_ = 0.7 nM) or recombinant LgBiT at saturating concentrations, establishing the maximum potential signal (**fig. S1B**). All protein designs exhibited robust signal enhancement when SmBiT^ultra^ or recombinant LgBiT was added, but high background luminescence was observed in samples with SmBiT^high^. The largest signal enhancements were achieved with SmBiT^low^, due to the reduced background complementation observed with this peptide (**fig. S1C**). Additional repeats of SmBiT did not yield greater light output, possibly due to insufficient spacing between sequential peptides to support LgBiT binding. We therefore moved forward with MCP-SmBiT^low^/PCP-LgBiT, the lantern combination that provided highest dynamic range.

We first asked whether the MCP-SmBiT^low^/PCP-LgBiT lantern could detect the previously reported MS2-PP7 bait, comprising an unstructured 19-nucleotide strand joining the two aptamers (**Fig. 1C**) (*14*). Upon RNA bait transcription, an approximate 17-fold signal increase was observed (**Fig. 1D**). We attributed the relatively low signal enhancement to the RNA bait structure. Secondary structure modeling with Vienna RNAfold (*21*) and Forna (*22*), revealed that the 19-nucleotide linker was likely flexible with the potential to sample non-productive conformations. While an unstructured RNA bait could aid lantern binding, we surmised that extensive flexibility could potentially hinder NanoBiT complementation and thus diminish signal-to-noise ratios.

We hypothesized that increasing the rigidity of the RNA bait would favor lantern assembly and photon production. We thus redesigned the RNA bait to fix the spacing and orientation of the MS2 and PP7 sequences. We locked the aptamers into desired conformations, as part of a four-way junction with an established Ni-NTA-binding aptamer (**Fig. 1E and fig. S2A**) (*23*). The RNA baits were further engineered to contain varying numbers of nucleotides between the MS2 and PP7 aptamer domains (M-X-P; where X is the number of nucleotides in the spacer). This panel enabled us to not only sample the spacing between the aptamer domains (∼3 nm, according to secondary structure predictions) but also the helical phase of the aptamers respective to one another (up to ∼360º). When the RNAs were present with the lantern fragments, we observed that a relatively short linker (M-3-P) could yield significantly higher luminescence, with up to a 330-fold increase over a no-RNA control (**Fig. 1E**). In the case of M-11-P, signal was abolished. We attributed this result to the phase angle of approximately 180º (one half of a helical turn) from M-3-P, preventing luciferase complementation. Importantly, signal was restored with the MS2 and PP7 aptamers brought back into phase (∼360º) and a full helical turn apart (M-21-P). Signal still decreased because the overall length increased and likely prevented effective assembly of the luciferase parts. Photon production was also highly dependent on the concentrations of the RNA bait and the RNA lantern (**fig. S2B**) and RNA bait integrity. In experiments using singular MS2 and PP7 aptamers, or M-3-P^mut^ that does not bind the PCP protein (**fig. S3**) (*24*), no appreciable signal over background was observed (**Fig. 1E and fig. S3C**).

Due to the improved luminescence achieved through modulating the distance and phase angle, we also examined the length and relative orientation of the individual MS2 and PP7 aptamers. We took the optimal M-3-P RNA bait and added or removed base-pairs (bp) to either the MS2 or PP7 stem (**Fig. 1F and fig. S4A**). No significant improvements in photon output were observed among the suite of RNA baits, suggesting that the length and orientation of the two aptamers were already optimal. We further confirmed that the lack of improvement was not a result of template DNA ratios (**fig. S4B**). Although the orientation of the aptamers had minimal effect on complementation, we found that the positioning of the aptamers was critical. When the placement of MS2 and PP7 were inverted, luciferase complementation was not observed (**fig. S5**).

We further examined the effect of protein linker length on RNA lantern assembly. Constructs were designed with additional glycine-serine (G_4_S) units inserted between the RNA-binding proteins and NanoBiT segments (**fig. S6A**). These constructs were then tested with the M-3-P, M-7-P, and M-11-P RNA baits (**fig. S6B**). The largest light outputs were achieved with the M-3-P RNA bait, for all the lanterns tested. Interestingly, only minimal signal enhancements (∼1.5 fold) were observed with the additional G_4_S units compared to the original lanterns both in vitro and (**fig. S6, C, D and E**).

The RNA sensing capabilities of the designer bait and lanterns were biochemically validated *in vitro* (**Fig. 2A**). Cells were engineered to stably express the lantern components, and lysates were then titrated with purified bait. As shown in Figures 2B-C, an RNA-dependent “hook effect” was observed (**Fig. 2, B and C**). Lantern complex formation was limited at low RNA concentrations, as expected. More lantern complementation was observed with increasing RNA bait, but the amount falls off at high RNA concentrations. This latter outcome is explained by MCP and PCP proteins binding distinct RNA bait transcripts, so that the luciferase bits are not brought together to assemble active lantern complexes.

**Fig. 2.**
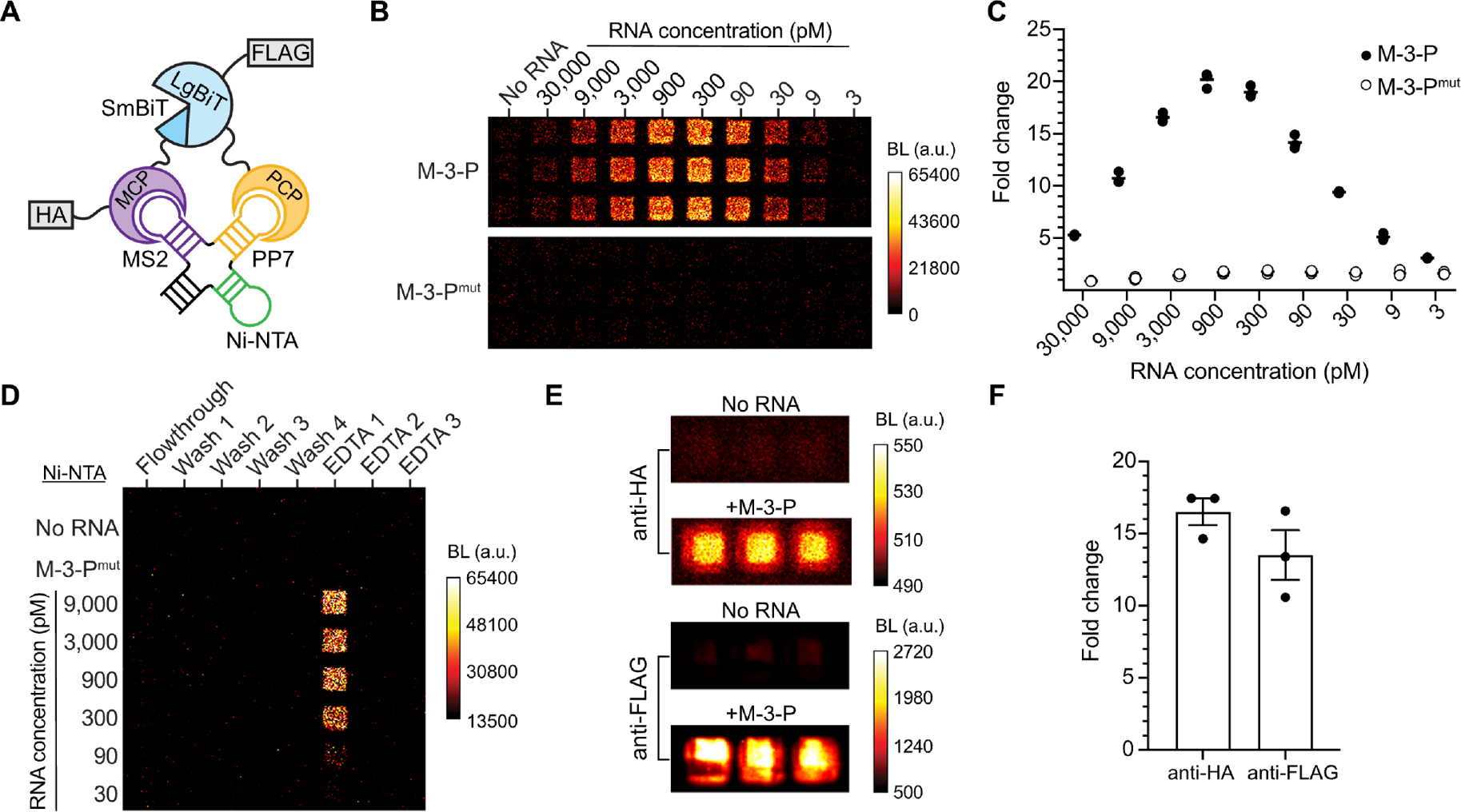
Biochemical validation of RNA lanterns. (**A**) The designer RNA bait comprises a 4-way-junction of MS2, PP7, and Ni-NTA aptamers. This unit assembles the RNA lantern components (MCP-SmBiT and PCP-LgBiT, fused to HA and FLAG epitopes, respectively). (**B**) Bioluminescent output from RNA bait (M-3-P) or inactive mutant (M-3-P^mut^) combined with lysate from cells stably expressing the RNA lantern. (**C**) Fold change in bioluminescent signal over no-RNA controls. Each bar represents the median of *n* = 3 replicates with dots showing data from individual replicates. (**D**) Affinity purification of the RNA lantern complex using the Ni-NTA aptamer. Various concentrations of RNA bait were used, and captured complexes were eluted using metal chelator (EDTA). (**E**) Bioluminescent output of lantern complexes captured using anti-HA or anti-FLAG conjugate-agarose beads. (**F**) Fold change in signal from lantern complexes retrieved in the presence of M-3-P bait versus no RNA from (E). Error bars represent the standard error mean (SEM) of *n* = 3 replicates. BL, bioluminescence. a.u., arbitrary units.

The unique design of the RNA bait enabled direct interrogation of lantern binding and assembly. The Ni-NTA aptamer was used to retrieve the RNA bait-RNA lantern complex on resin: as shown in Figure 2D, the assembled NanoBiT enzyme was exclusively associated with the RNA bait (**Fig. 2D**). Pulldowns of active complexes using the HA and FLAG tags built into the RNA lantern components further confirmed the RNA-dependent assembly of the active luciferase (**Fig. 2, E and F**). We anticipate that the retrieval of targeted transcripts and associated biomolecules post-imaging will facilitate the often-critical follow-up analyses of RNA interactions.

### RNA imaging platform is ultrasensitive and modular

Previous applications of MS2-PP7 for fluorescence imaging have required at least 12 copies of the aptamers to achieve adequate signal-to noise ratios (MP_12X,_ **Fig. 3A**) (*2, 15, 25, 26*). In many cases, these constructs comprise an RNA bait that is at least 780 nucleotides long, a size that may impede the natural localization and behavior of the RNA under study (*27-29*). The M-3-P bait (69 nucleotides) requires only a single copy of each aptamer to produce detectable signal (**Fig. 3B**). When compared head-to-head with bait aptamers at equimolar concentration (M-3-P versus one twelfth RNA concentration of MP_12X_), the structured bait produced 10-times more signal. Even when MP_12X_ was introduced at the same RNA concentration and the aptamers were 12-times more concentrated, the structured M-3-P bait outperformed the flexible design (**Fig. 3B**). These results suggest that the compact M-3-P is a more effective RNA bait, providing higher signal compared to the multimerized and extensively used MS2-PP7 tag.

**Fig. 3.**
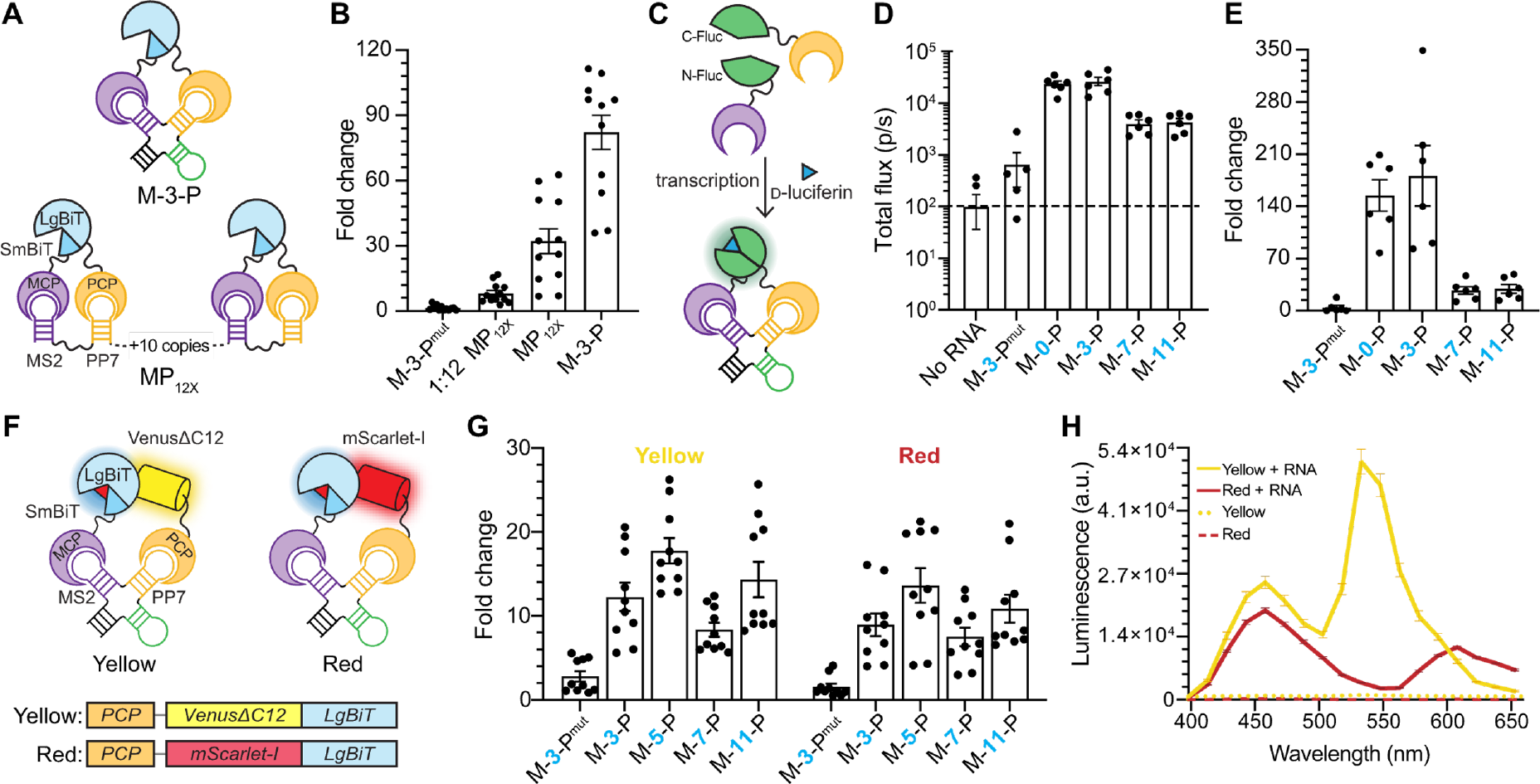
Robustness and modularity of structured RNA baits. (**A**) Comparison of M-3-P to a 12-copy unstructured RNA bait. Top: Schematic of the structured M-3-P RNA bait (single copy). Bottom: Schematic of a flexible RNA bait comprising 12 copies of the MS2 and PP7 aptamers (MP_12X_). (**B**) MP_12x_ was evaluated against M-3-P at equimolar RNA concentrations (1 nM of template DNA) or at 1/12 the concentration (83 pM of template DNA). Fold change over no-RNA bait samples is plotted. (**C**) RNA bait assembles other lanterns. Schematic of MCP-PCP probes fused to split firefly luciferase (Fluc). MCP is fused to the N-terminal half of Fluc and PCP is fused to the C-terminal half. RNA transcription followed by D-luciferin treatment enables photon production and visualization. (**D**) The Fluc lantern was assessed with a panel of RNA baits. Total light output for *n* = 6 replicates is shown. Only 2 replicates are shown for the no-RNA bait sample; the others were below the limit of detection represented by the electronic noise of the EMCCD camera (dashed line). (**E**) Graph of fold change in signal over no-RNA bait samples from (D). (**F**) Schematic of BRET-based RNA lanterns. (**G**) Luminescence fold change measurements for yellow and red RNA lanterns across a panel of RNA baits. Fold change calculated over no-RNA controls. (**H**) Emission spectra for BRET-based RNA lanterns. Error bars represent the standard error of the mean (SEM) for *n* = 12 replicates (**Fig. 3B**), *n* = 6 replicates (**Figs. 3D** and **3E**), *n* = 10 replicates (**Fig. 3G**), and *n =* 3 replicates (**Fig. 3H**). a.u., arbitrary units.

The designer M-3-P bait can also facilitate complementation of other split reporters. When MCP and PCP were fused to split fragments of firefly luciferase (Fluc; **Fig. 3C**), the resulting RNA lantern yielded a large increase in signal over background (**Fig. 3, D and E**). The photon flux of the assembled Fluc is lower than NanoBiT, but the undetectable background in absence of RNA may be desirable in specific applications. M-3-P also enabled the assembly of bioluminescence resonance energy transfer (BRET) variants of NanoLuc. These probes comprise luciferase fluorescent protein fusions that produce red-shifted light upon complementation. Such wavelengths can provide more sensitive readouts in vivo because they are less absorbed and scattered by tissue. Two BRET-based RNA lanterns were developed using yellow (VenusΔC12) (*9*) and red (mScarlet-I) (*30*) fluorescent proteins fused to the C-terminus of PP7 and the N-terminus of LgBiT (**Fig. 3F**). Assembly of the yellow and red RNA lanterns was assessed by measuring photon production in the presence of rigidified RNA baits. As shown in Figure 3G, M-5-P provided the highest degree of signal turn-on (**Fig. 3G**). The larger bait provides more space between the bulkier lantern fragments in these cases, likely enabling more effective complementation. The BRET-bases probes also exhibited different colors of light output (**Fig. 3H**). BRET efficiency was higher for the yellow than the red lantern due to the larger spectral overlap between NanoBiT and VenusΔC12 compared to mScarlet-I (*9, 31*). Collectively, these demonstrate that structured RNA baits can productively assemble diverse split proteins.

### Imaging RNAs from the micro-to-macro scale

The robustness of the bait design enabled facile imaging of RNA both *in cellulo* and *in vivo*. As an initial test, we developed a model system using an mRNA encoding green fluorescent protein (GFP) with the M-3-P RNA bait placed in the 3′ untranslated region (UTR) (**Fig. 4A**). GFP fluorescence would thus report on bait expression. Cell lines stably expressing RNA lanterns were generated and transfected with constructs encoding M-3-P–tagged *GFP* or *GFP* only. GFP fluorescence was observed in both cases, but bioluminescence (from RNA lantern assembly) was only observed in cells transfected with M-3-P–tagged *GFP* (**Fig. 4, B and C, and fig. S7**). Similar bioluminescence turn-on was observed using an analogous model transcript (**fig. S6, D and E**). Both studies illustrate the utility of the structured bait for RNA lantern assembly and transcript visualization.

**Fig. 4.**
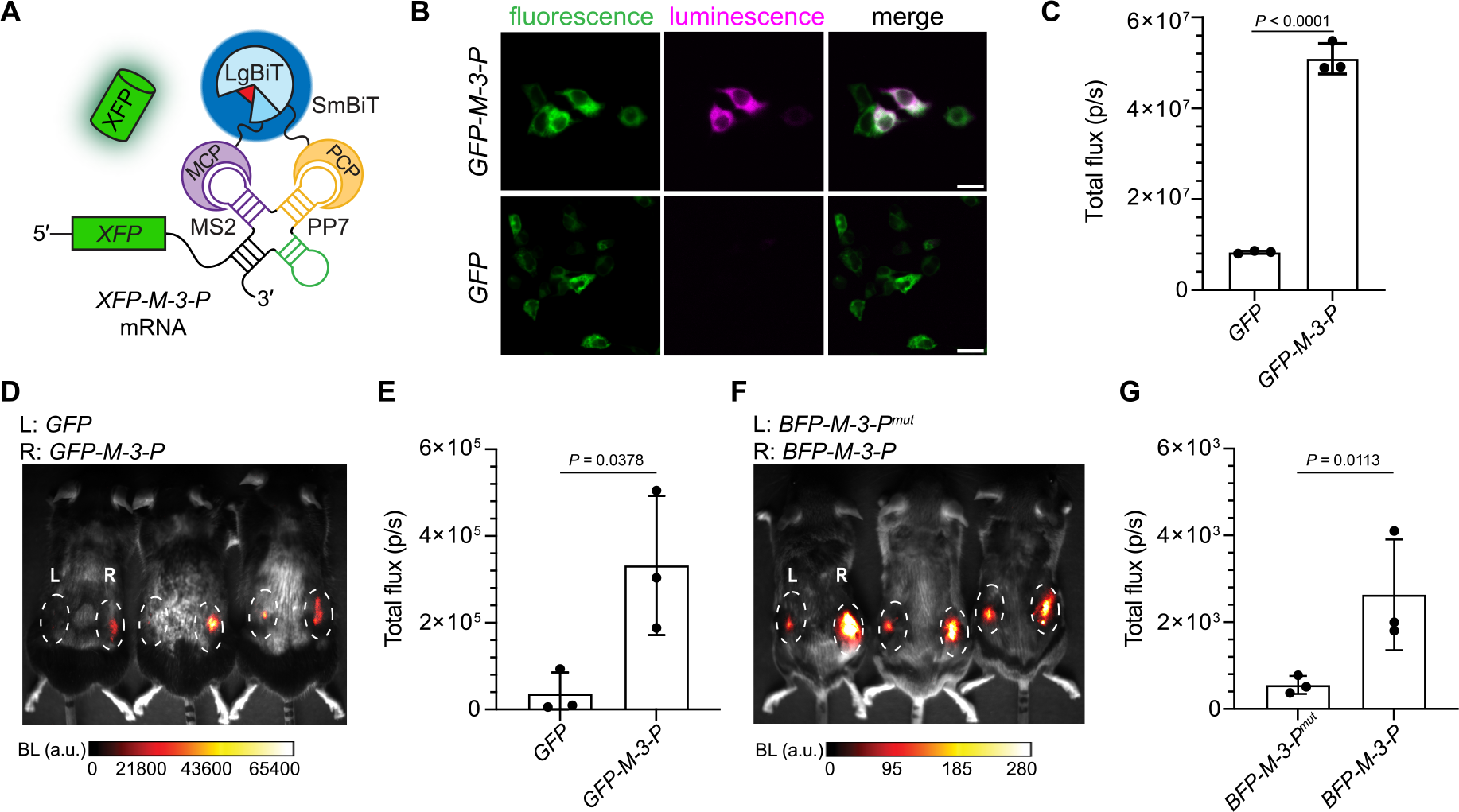
Imaging in mammalian cells and live mice with RNA lanterns. (**A**) Schematic of model mRNA encoding a fluorescent protein. The sequence was engineered with M-3-P in the 3′ UTR. Transcription of *XFP-M-3-P* mRNA results in lantern assembly and light emission (blue glow). XFP production can be further analyzed via fluorescence (green glow). (**B**) HEK293T cells expressing the RNA lantern were transiently transfected with DNA encoding *GFP–M-3-P* (Pearson’s r = 0.84) or *GFP* alone (Pearson’s r = 0.03). Luminescence was observed exclusively in cells containing mRNAs with the M-3-P bait. Scale bar = 20 µm. (**C**) Bulk measurement of photon flux from lantern-expressing cells transfected with DNA encoding *GFP* or *GFP–M-3-P*. (**D**) Mice implanted with cells expressing RNA lanterns and *GFP* alone (left flank) or *GFP-M-3-P* (right flank) and imaged with luciferin. (**E**) Photon flux from mice with implanted cells expressing *GFP(±)M-3-P* and RNA lanterns shown in (D). (**F**) Mice implanted with cells expressing RNA lanterns and *BFP-M-3-P*^*mut*^ (left flank) or *BFP-M-3-P* (right flank). (**G**) Photon flux from mice with implanted cells expressing either *BFP-M-3-P* or *BFP-M-3-P*^*mut*^ and RNA lanterns. Error bars represent standard deviation (SD) for *n* = 3 replicates (Fig. 4C, 4E, and 4G). *P*-values (95% confidence interval) were determined by unpaired, two-tailed *t*-test. BL, bioluminescence. a.u., arbitrary units.

Finally, as a proof of concept, we imaged RNAs in subcutaneous models *in vivo*. HEK293T cells were engineered to express RNA lanterns and mRNAs encoding either GFP or BFP with variable 3′ UTRs: M-3-P, M-3-P^mut^, or no bait (**Fig. 4A**). The cells were incubated with luciferin, implanted in mice dorsal flanks, and imaged. Increased luminescence was observed from cells expressing RNA lanterns and *GFP*-*M-3-P* (right flank) compared to cells lacking the M-3-P bait (left flank (**Fig. 4, D and E**). Similar increases in luminescence were observed from cells expressing *BFP*-*M-3-P*, compared to control transcripts in the presence of the lantern components (**Fig. 4, F and G**). These data demonstrate that lanterns can be used for RNA-dependent imaging in tissue, setting the stage for real-time detection of gene expression in live animals.

## Discussion

RNA dynamics have been historically visualized in living systems with fluorescent probes. Such methods require optically permissive platforms such as transparent model organisms or surgically implanted windows. The required excitation light can further induce high background signals in tissue and is often limited in depth. Bioluminescent probes (luciferases) overcome some of these limitations. Luciferase reporters use enzymatic reactions—instead of excitation light—to generate photons, which produces far less background signal and can be advantageous for serial imaging in tissue and whole animals.

To capitalize on the sensitivity and dynamic range of bioluminescence, we developed a genetically encoded split luciferase-based platform for visualizing RNA transcripts. The RNA lanterns combine the well-known NanoBiT system and MS2/PP7 platform for transcript tracking. We systematically optimized the components to achieve sensitive imaging. Substantial improvements in signal production were achieved by modulating the spacing and phase angle of the aptamer components. The optimized design significantly decreases the size of previously reported RNA-protein complexes for imaging. Only a single copy of the RNA bait was necessary for imaging target transcripts in cells and whole organisms.

The RNA bait and lanterns also comprise unique features for a range of applications. Both the proteins and bait are equipped with affinity tags for retrieval and downstream analyses of target transcripts and interacting biomolecules. Structured RNA baits can productively assemble diverse split proteins, including Fluc and various BRET reporters. Such modularity expands the color palette of RNA lanterns and sets the stage for even deeper tissue imaging and multiplexed assays. Given our finding that the BRET probes were more efficiently assembled with longer RNA baits, though, care should be taken to optimize each lantern and tag combination.

We anticipate that RNA lanterns will enable RNA dynamics to be visualized in vitro and in vivo. The probes will be particularly useful for serial imaging of transcripts, where localization and expression are difficult to examine over time. While we focused on visualizing model transcripts with RNA bait incorporated into the 3′ UTR, the short tag can easily be applied to other transcripts through genetic manipulation. Further tuning of both the RNA lantern and tag will expand the number of transcripts that can be visualized in tandem. Such multiplexed studies will paint a more complete picture of RNA biology *in vivo*.

## Acknowledgments

We thank Dr. Lorenzo Scipioni and Prof. Michelle Digman (UCI) for help with optical imaging experiments. We also thank members of the Luptak, Steward, and Prescher laboratories for helpful discussions.

## Funding

W.M. Keck Foundation: OS, AL, JAP

NASA ICAR 80NSSC21K0596: AL

The Paul G. Allen Frontiers Group: JAP

National Science Foundation 1804220: AL

National Institute of Health grant R01CA229696: CCC

## Author contributions

Conceptualization: CCC, OS, AL, JAP

Methodology: KKN, KHC, LPH, AL, JAP

Investigation: KKN, KHC, LPH, CETC, EBF, CC, MM, CCC, AAB, ER-M, ADK

Visualization: KKN, LPH, KHC

Writing – original draft: KKN, LPH, KHC, AL, JAP

Writing – review and editing: all authors

Funding acquisition: OS, SL, JAP

## Competing interests

AL, JAP, OS, KKN, KHC, LPH, and CCC are on a provisional patent application, filed through UCI, based on the results described here.

## Data and materials availability

All data are presented in the main text and supplementary materials. Plasmid DNAs are available upon request.

## Notes

### Competing Interest Statement

A provisional patent application has been filed through UCI based on the results described here.

### Summary of Updates

Additional results and discussion have been added; Supplemental files updated.

## References and Notes

1. P. A. Sharp, The Centrality of RNA. Cell 136, 577–580 (2009).

2. D. R. Larson, D. Zenklusen, B. Wu, J. A. Chao, R. H. Singer, Real-Time Observation of Transcription Initiation and Elongation on an Endogenous Yeast Gene. Science 332, 475–478 (2011).

3. J. M. Halstead et al., An RNA biosensor for imaging the first round of translation from single cells to living animals. Science 347, 1367–1671 (2015).

4. J. S. Paige, K. Y. Wu, S. R. Jaffrey, RNA mimics of green fluorescent protein. Science 333, 642–646 (2011).

5. S. Das, H. C. Moon, R. H. Singer, H. Y. Park, A transgenic mouse for imaging activity-dependent dynamics of endogenous Arc mRNA in live neurons. Sci Adv 4, eaar3448 (2018).

6. B. H. Lee et al., Real-time visualization of mRNA synthesis during memory formation in live mice. Proceedings of the National Academy of Sciences 119, e2117076119 (2022).

7. L. Jiang et al., Large Stokes shift fluorescent RNAs for dual-emission fluorescence and bioluminescence imaging in live cells. Nature Methods 20, 1563–1572 (2023).

8. M. P. Hall et al., Engineered Luciferase Reporter from a Deep Sea Shrimp Utilizing a Novel Imidazopyrazinone Substrate. ACS Chemical Biology 7, 1848–1857 (2012).

9. K. Suzuki et al., Five colour variants of bright luminescent protein for real-time multicolour bioimaging. Nature Communications 7, 13718 (2016).

10. F. X. Schaub et al., Fluorophore-NanoLuc BRET Reporters Enable Sensitive In Vivo Optical Imaging and Flow Cytometry for Monitoring Tumorigenesis. Cancer Res 75, 5023–5033 (2015).

11. H.-W. Yeh et al., ATP-Independent Bioluminescent Reporter Variants To Improve in Vivo Imaging. ACS Chemical Biology 14, 959–965 (2019).

12. M. Eguchi, H. Yoshimura, Y. Ueda, T. Ozawa, Split Luciferase-Fragment Reconstitution for Unveiling RNA Localization and Dynamics in Live Cells. ACS Sensors, (2023).

13. F. Lim, T. P. Downey, D. S. Peabody, Translational repression and specific RNA binding by the coat protein of the Pseudomonas phage PP7. J Biol Chem 276, 22507–22513 (2001).

14. A. Bernardi, P.-F. Spahr, Nucleotide Sequence at the Binding Site for Coat Protein on RNA of Bacteriophage R17. Proceedings of the National Academy of Sciences 69, 3033–3037 (1972).

15. B. Wu, J. Chen, R. H. Singer, Background free imaging of single mRNAs in live cells using split fluorescent proteins. Scientific Reports 4, 3615 (2014).

16. A. S. Dixon et al., NanoLuc Complementation Reporter Optimized for Accurate Measurement of Protein Interactions in Cells. ACS Chemical Biology 11, 400–408 (2016).

17. H. W. Yeh, H. W. Ai, Development and Applications of Bioluminescent and Chemiluminescent Reporters and Biosensors. Annu Rev Anal Chem (Palo Alto Calif) 12, 129–150 (2019).

18. C. R. Bodle, M. P. Hayes, J. B. O’Brien, D. L. Roman, Development of a bimolecular luminescence complementation assay for RGS: G protein interactions in cells. Anal Biochem 522, 10–17 (2017).

19. T. Machleidt et al., NanoBRET—A Novel BRET Platform for the Analysis of Protein– Protein Interactions. ACS Chemical Biology 10, 1797–1804 (2015).

20. T. D. Goddard et al., UCSF ChimeraX: Meeting modern challenges in visualization and analysis. Protein Sci 27, 14–25 (2018).

21. R. Lorenz et al., ViennaRNA Package 2.0. Algorithms for Molecular Biology 6, 26 (2011).

22. P. Kerpedjiev, S. Hammer, I. L. Hofacker, Forna (force-directed RNA): Simple and effective online RNA secondary structure diagrams. Bioinformatics 31, 3377–3379 (2015).

23. F. Chizzolini et al., Large Phenotypic Enhancement of Structured Random RNA Pools. Journal of the American Chemical Society 142, 1941–1951 (2020).

24. F. Lim, D. S. Peabody, RNA recognition site of PP7 coat protein. Nucleic acids research 30, 4138–4144 (2002).

25. T. Lionnet et al., A transgenic mouse for in vivo detection of endogenous labeled mRNA. Nat Methods 8, 165–170 (2011).

26. E. Tutucci et al., An improved MS2 system for accurate reporting of the mRNA life cycle. Nat Methods 15, 81–89 (2018).

27. J. F. Garcia, R. Parker, MS2 coat proteins bound to yeast mRNAs block 5′ to 3′ degradation and trap mRNA decay products: implications for the localization of mRNAs by MS2-MCP system. RNA 21, 1393–1395 (2015).

28. S. Heinrich, C. L. Sidler, C. M. Azzalin, K. Weis, Stem–loop RNA labeling can affect nuclear and cytoplasmic mRNA processing. RNA 23, 134–141 (2017).

29. J. F. Garcia, R. Parker, Ubiquitous accumulation of 3′ mRNA decay fragments in Saccharomyces cerevisiae mRNAs with chromosomally integrated MS2 arrays. RNA 22, 657–659 (2016).

30. D. S. Bindels et al., mScarlet: a bright monomeric red fluorescent protein for cellular imaging. Nature Methods 14, 53–56 (2017).

31. K. K. Ng, J. A. Prescher, Generalized Bioluminescent Platform To Observe and Track Cellular Interactions. Bioconjugate Chemistry 33, 1876–1884 (2022).

32. K. Valegârd et al., The three-dimensional structures of two complexes between recombinant MS2 capsids and RNA operator fragments reveal sequence-specific protein-RNA interactions. J Mol Biol 270, 724–738 (1997).

33. J. A. Chao, Y. Patskovsky, S. C. Almo, R. H. Singer, Structural basis for the coevolution of a viral RNA-protein complex. Nat Struct Mol Biol 15, 103–105 (2008).

34. D. G. Gibson et al., Enzymatic assembly of DNA molecules up to several hundred kilobases. Nature Methods 6, 343–345 (2009).

35. S. Bolte, F. P. CordeliÉRes, A guided tour into subcellular colocalization analysis in light microscopy. Journal of Microscopy 224, 213–232 (2006).

