## Supplementary Materials for "A modular platform for bioluminescent RNA tracking"

### Materials and Methods

General information

Q5 DNA polymerase, restriction enzymes, and all buffers were purchased from New England Biolabs. dNTPs were purchased from Thermo Fisher Scientific. Luria-Bertani medium (LB) was purchased from Genesee Scientific. All plasmids and primer stocks were stored at -20 °C unless otherwise noted. Primers were purchased from Integrated DNA Technologies and plasmids were sequenced by Azenta Life Sciences. Sequencing traces were analyzed using Benchling.

General cloning methods

Polymerase chain reaction (PCR) was used to prepare genes of interest, and products were analyzed by gel electrophoresis. Products were excised and purified. Amplified genes were ligated into destination vectors via Gibson assembly (*34*). All PCR reactions were performed in a BioRad C3000 thermocycler using the following conditions: 1) 95 °C for 3 min, 2) 95 °C for 30 s, 3) –1.2°C per cycle starting at 72 °C for 30 s, 4) 72 °C for 30 sec, repeat steps 2–4 ten times, 5) 95 °C for 3 min, 6) 95 °C for 30 s, 7) 60 °C for 30 s, 8) 72 °C for 2 min repeat steps 6–8 twenty times, then 72 °C for 5 min, and hold at 12 °C until retrieval from the thermocycler. Gibson assembly conditions were: 50 °C for 60 min and held at 12 °C until retrieval from the thermocycler. Ligated plasmids were transformed into TOP10 *E. coli* cells using the heat shock method. After incubation at 37 °C for 18–24 h, colonies were picked and expanded overnight in 5 mL LB broth supplemented with ampicillin (100 µg/mL) or kanamycin (100 µg/mL). DNA was extracted from colonies using a Zymo Research Plasmid Mini-prep Kit. DNA was subjected to restriction enzyme digestion to confirm gene insertion. Positive hits were further sequenced. See Table S1 for list of constructs.

General cell culture methods

HEK293T cells (HEK, ATCC) and stable cells lines derived from HEK293T cells were cultured in complete media: DMEM (Corning) containing 10% (v/v) fetal bovine serum (FBS, Life Technologies), penicillin (100 U/mL), and streptomycin (100 µg/mL, Gibco). Cell lines stably expressing RNA lantern and linker designs were generated via lentiviral transduction. Transduced cells were further cultured with puromycin (20 µg/mL) to preserve gene incorporation. Cells were incubated at 37 °C in a 5% $CO_2$ humidified chamber. Cells were serially passaged using trypsin (0.25 % in HBSS, Gibco).

Protein expression and purification

LgBiT was encoded in a pCold vector. LgBiT was expressed in *E. coli* BL21 cells grown in LB medium (1 L). Expression was induced at an optical density (OD600) of ~0.6 by addition of 0.5 mM isopropyl-β-D-thiogalactopyranoside (IPTG), followed by incubation at 37 °C for 4 h. Cells were harvested by centrifugation (4,000 x*g*, 10 min, 4 °C). Cells were then resuspended in lysis buffer (30 mL, 50 mM Tris HCl, 150 mM NaCl, 0.5% Tween-20, 1 mM phenylmethylsulfonyl fluoride [PMSF], pH 7.4). Cells were sonicated (Qsonica) at 40% amplitude, at 2 sec on 2 sec off intervals for 15 min. Cell debris was removed through centrifugation (10,000 x*g*, 1 h, 4 °C). Proteins were purified by Ni-NTA affinity chromatography. The column was washed with wash buffer (20 mM imidazole, 50 mM $NaPO_4$, pH 7.4). Protein was eluted from column with elution buffer (200 mM imidazole, 50 mM $NaPO_4$, pH 7.4). Protein was dialyzed overnight at 4 °C into phosphate buffer (50 mM $NaPO_4$, pH 7.4). Protein was concentrated to ~500 µL using Amicon

Ultra-15 Centrifugal Filter Units (Merck Millipore MWCO 3 kDa). The concentration of proteins was determined using JASCO V730 UV-vis spectrophotometer at 280 nm. SDS-PAGE was also performed to verify purity, and gels were stained with Coomassie R-250.

*In vitro* transcription

RNA was transcribed *in vitro* in a buffer containing 50 mM Tris-HCl pH 8, 2 mM spermidine, 0.01% Triton X-100, 2 mM rNTPs (each), 20 mM $MgCl_2$, 10 mM dithiothreitol (DTT), and recombinant T7 RNA polymerase. Transcription reactions were quenched with 30 mM EDTA and subsequently ethanol-precipitated with coprecipitant (Invitrogen GlycoBlue, AM9516) prior to purification via denaturing PAGE. Purified RNAs were eluted from gel pieces into 300 mM KCl and 0.1 mM EDTA for 3 h. Eluted RNAs were ethanol-precipitated with coprecipitant, and pellets were washed with 70% ethanol, dried, and resuspended in 10 mM Tris-HCl pH 7.5, 0.1 mM EDTA, and 0.001% Triton X-100 (TET). RNA concentrations were determined by UV-vis absorption spectroscopy (Thermo Scientific NanoDrop 2000, ND-2000).

*In vitro* transcription/translation (IVTT) in rabbit reticulocyte lysate

A rabbit reticulocyte lysate *in vitro* translation kit (Promega, L4960) was coupled to *in vitro* transcription with the supplementation of $MgCl_2$, rNTPs, and T7 RNA polymerase. Plasmid DNAs were 3′ linearized for run-off transcription. RNA scaffolds were prepared via primer elongation of synthetic DNA oligos (Integrated DNA Technologies). Linearized plasmid DNAs and RNA bait DNAs were kit purified prior to IVTT (DNA Clean and Concentrator; Zymo Research, D4004). All DNAs were eluted into Tris-EDTA (TE; 10 mM Tris-HCl and 0.1 mM EDTA, pH 7.5) and controls were supplemented with an equal volume of TE buffer.

IVTT reactions were prepared on ice with 70% v/v lysate, 0.02 mM amino acid mix, rNTPs (0.35 mM GTP and 0.15 mM each ATP/CTP/UTP), 1 mM $MgCl_2$, linearized plasmid DNA, RNA bait template DNA, 2 U/10 µL RNase inhibitor (Invitrogen, AM2694), and 0.5 µL/10 µL reaction of recombinant T7 RNA polymerase (house preparation). NanoBiT experiments were set up with 10 µL volumes and 1 nM of linearized plasmid DNA. Split firefly luciferase experiments were performed with 25 µL volumes and 3 nM of linearized plasmid DNA. RNA bait DNA templates were varied at specified ratios. IVTT reactions were incubated at 34 °C for 10 min and 30 °C for 140 min.

Luciferin substrates were supplied prior to imaging. For split NanoBiT RNA lantern experiments, furimazine was added to each reaction at 20 µM, and samples were incubated at room temperature for 5 min before imaging. For split firefly luciferase RNA lantern experiments, D-luciferin (100 µM) and ATP (1 mM) were added to each reaction. Samples were then incubated at room temperature for 5 min before image acquisition. Plates were imaged in a dark, light-proof chamber using an IVIS Lumina (PerkinElmer) CCD camera chilled to −90 °C. The stage was kept at 37 °C during imaging and the camera was controlled using Living Image software. Exposure times were set to 1 min and binning levels were set to medium. Regions of interest were selected for quantification and total flux values were analyzed using Living Image software. All data was exported to Microsoft Excel or Prism (GraphPad) for further analysis.

*In cellulo* linker optimization

Stable cell lines expressing variant RNA lanterns were plated in clear 12-well plates and incubated overnight. Cell lines were then transfected with 1 μL Lipofectamine 3000 (Invitrogen, L3000001), 2 μL P3000 Reagent (Invitrogen, L3000001), and 500 ng of the BFP-M-3-P plasmid. After 24 h incubation, cells were lifted and counted with a Countess II (Invitrogen). 50,000 transfected or non-transfected cells were plated in triplicate into black 96-well plates (Greiner Bio-One). Furimazine (20 μM) was added to each sample. Plates were imaged in a dark, light-proof chamber using an IVIS Lumina (PerkinElmer) CCD camera chilled to −90 °C. The stage was kept at 37 °C during imaging and the camera was controlled using Living Image software. Exposure times were set to 1 min and binning levels were set to medium. Regions of interest were selected for quantification and total flux values were analyzed using Living Image software. All data was exported to Microsoft Excel or Prism (GraphPad) for further analysis. Remaining cells were analyzed on a Novocyte flow cytometer (ACEA BioSciences) for BFP expression.

General flow cytometry methods

Cells were treated with trypsin for 5 min at 37 °C and then neutralized with complete media. Cells were transferred to Eppendorf tubes and pelleted (500 x*g*, 5 min) using a tabletop centrifuge (Thermo Fisher Sorvall Legend Micro 17). The resulting supernatants were discarded, and cells were washed with PBS (2 × 400 μL). Cells were analyzed for XFP expression on a Novocyte flow cytometer. Live cells were gated, and singlet cells were further gated. For each sample, 10,000 events were collected on the "singlet cell" gate.

HEK293T lysate preparation

HEK293T cells stably expressing RNA lanterns (~$10^6$) were pelleted at 500 x*g* for 10 min at 4 °C and washed three times with binding buffer (140 mM KCl, 10 mM NaCl, 20 mM Tris-HCl pH 7.5, and 5 mM $MgCl_2$). Cells were pelleted during each washing step at 500 x*g* (1 min, 4 °C) After the final wash, the cell pellet was resuspended in 600 μL binding buffer supplemented with 1% Tween-80, 1X protease inhibitor cocktail (Roche, 11697498001), and 5 μg RNase inhibitor. Cells were sonicated on ice (5 cycles of 10 s on, 10 s off; Branson CPX2800). Cell debris was then removed by centrifugation (12,000 x*g*, 4 °C, 15 min). The resulting supernatant was aliquoted and stored at –80 °C for future use as 2X lysate stocks.

RNA titration assay

Purified RNAs (10X stocks in TET buffer) were serially diluted. Binding reactions contained 12.5 μL of 2X cell lysate, 10 μL of 1X binding buffer, and 2.5 μL of 10X RNA. Reactions were incubated at room temperature on an orbital shaker for 30 minutes. After incubation, 1 μL of a 200 μM furimazine stock (Promega, Nano-Glo substrate) in 1X binding buffer was added to each reaction and then plated in a black, clear bottom 384-well plate and imaged with an Andor iXon Ultra 888 EMCCD camera (Oxford Instruments) equipped with an F 0.95 lens (Schneider) with an EM gain of 500. Images were acquired (5 s acquisition) over 10 min. Images were Z-stacked in ImageJ and densitometry was used to measure peak intensity values.

Ni-NTA pull-down

Triplicate reactions of selected RNA concentrations above, within, and below the curve (as determined by RNA titration assay) were combined (45 μL/each). Charged Ni-NTA agarose beads (~45 μL; Thermo Fisher, 88221) were washed three times in 90 μL 1X binding buffer in a spin-column (Corning, 8160) at 3000 x*g* for 30 s. The combined binding reactions were then incubated on the Ni-NTA agarose at room temperature for 30 min on an orbital shaker. After incubation, the columns were spun at 3000 x*g* for 30 s and the flowthrough was collected. Four 5-min washes were performed with 45 µL 1X binding buffer at room temperature on an orbital shaker. After each incubation, the columns were spun, and the flowthroughs were collected (washes 1-4). Columns were eluted three times with 1X binding buffer supplemented with 25 mM EDTA (20 mM final) and incubated for 5 min on an orbital shaker. Elution fractions were collected. Furimazine was added to the fractions (20 µM final) and the solutions were placed in a black, clear bottom 384-well plate. Images were acquired using an Andor iXon Ultra 888 EMCCD camera equipped with an F 0.95 lens, with an EM gain of 500. Images were acquired (10 s acquisitions) over 15 min. Final images were Z-stacked, and densitometry (ImageJ) was used to determine fold change over background and no RNA controls. Final images were overlaid with brightfield photos. Plots were produced using GraphPad Prism.

HA and FLAG pull-down

Reticulocyte lysate IVTT reactions (67.5 µL) were set up as previously described. To each reaction 100 nM M-3-P RNA (7.5 µL) or 1X binding buffer (7.5 µL) was added prior to incubation. Reactions were incubated at 34 °C for 10 min and 30 °C for 140 min. Pierce magnetic anti-HA beads (45 µL) and anti-FLAG beads (1.5 µL) were normalized for binding capacity. Two sets of each bead were washed three times in 1X binding buffer ten-times their initial volume (450 µL and 15 µL, respectively; 5 min with agitation). To each tube, 75 µL of IVTT reactions were added and incubated with agitation for 60 min at room temperature. The supernatant was removed and stored separately. After incubation, beads were washed three times in 1X binding buffer (75 µL, 5 min with agitation) and each wash was stored. Beads were resuspended in 30 µL 1X binding buffer and 20 µM furimazine, then plated in triplicate in a black, clear bottom 384-well plate and imaged with an Andor iXon Ultra 888 EMCCD camera equipped with an F 0.95 lens, with an EM gain of 100. Images were acquired (5 s acquisition) over 15 min. Images were Z-stacked in ImageJ and densitometry was used to measure peak intensity values.

Bioluminescence microscopy

HEK293T cells ($5 \times 10^5$) stably expressing RNA lanterns were plated in 8-well Ibidi µ-Slides. After 24 h, the cells were transiently transfected with 100 ng of *GFP*-M-3-P plasmid or GFP plasmid using 0.3 µL Lipofectamine 3000 and 0.3 µL P3000 reagent. After 18 h, cell media was exchanged for phenol red-free DMEM (Gibco FluoroBrite DMEM, A1896701) supplemented with 20 µM furimazine. Live cell images were captured on an Olympus IX71 microscope equipped with an Andor iXon Ultra 888 EMCCD camera and a 40X oil objective (Olympus UPlanApo 40X/1.00 oil iris). Images were captured with an EM gain of 1000, 10 MHz horizontal read-out rate, 4.33 µs vertical clock speed, and an acquisition time of 180 s. The microscope stage was kept warm with a heating pad to maintain cell viability and to encourage enzyme turnover. Fluorescence images were captured immediately following luminescence imaging, using a blue LED light source (ThorLabs, Solis-470C). Fluorescent images were captured using the EMCCD's conventional mode with an acquisition time of 0.5 s. Images were processed (removed outliers and Z-stacked) with ImageJ and colocalization was determined with the JaCoP plug-in (*35*).

*In vivo* cell transplants

Mouse experiments were approved by the UC Irvine Animal Care and Use Committee (IACUC) in compliance with the National Institute of Health guidelines. Mice were maintained on a 12 h light/dark cycle at 25 °C. All procedures were done during the light portion of the cycle. Our studies were done using Rosa26$^{tdTomato}$ male mice obtained from the Jackson Laboratory (JAX: 007905).

Mice were anesthetized using isoflurane (1-2%) and were given subcutaneous dorsal injections of HEK293T cells ($1 \times 10^6$) expressing RNA lanterns and *GFP*, *GFP-M-3-P*, *BFP-M-3-P$^{mut}$*, or *BFP-M-3-P* suspended in sterile PBS solution. Cells were normalized for mean fluorescence intensity (MFI) of *XFP* using flow cytometry. Following normalization, cells were implanted into the dorsal posterior subcutaneous flank of each mouse. Cells expressing RNA lantern and controls, *GFP* or *BFP-M-3-P$^{mut}$*, were implanted on the left side, while cells expressing the RNA lantern and *GFP-M-3-P* or *BFP-M-3-P* were implanted on the right side, allowing each mouse to serve as its own control. Each animal received a 20 µM furimazine injection in 200 µL of implant.

Animals were imaged immediately following transplantation. For imaging, animals were anesthetized with an i.p. injection of ketamine (6.6 mg/mL) and xylazine (1.65 mg/mL) suspended in sterile PBS and placed on a warmed (37 °C) stage. Images were captured on an Andor iXon Ultra 888 EMCCD camera equipped with an F 0.95 lens. Images were acquired with an EM gain of 1000, an acquisition time of 300 s, and signal was captured as photons. Images were processed (removing outliers and integrated density) using ImageJ. Total flux (p/s) was determined based on a given region of interest (ROI) for each implantation. Background correction was accomplished by averaging ROIs containing no luminescence. Prism (GraphPad) was used to determine significant differences (unpaired, two-tailed *t*-test) between groups.

**Fig. S1.**

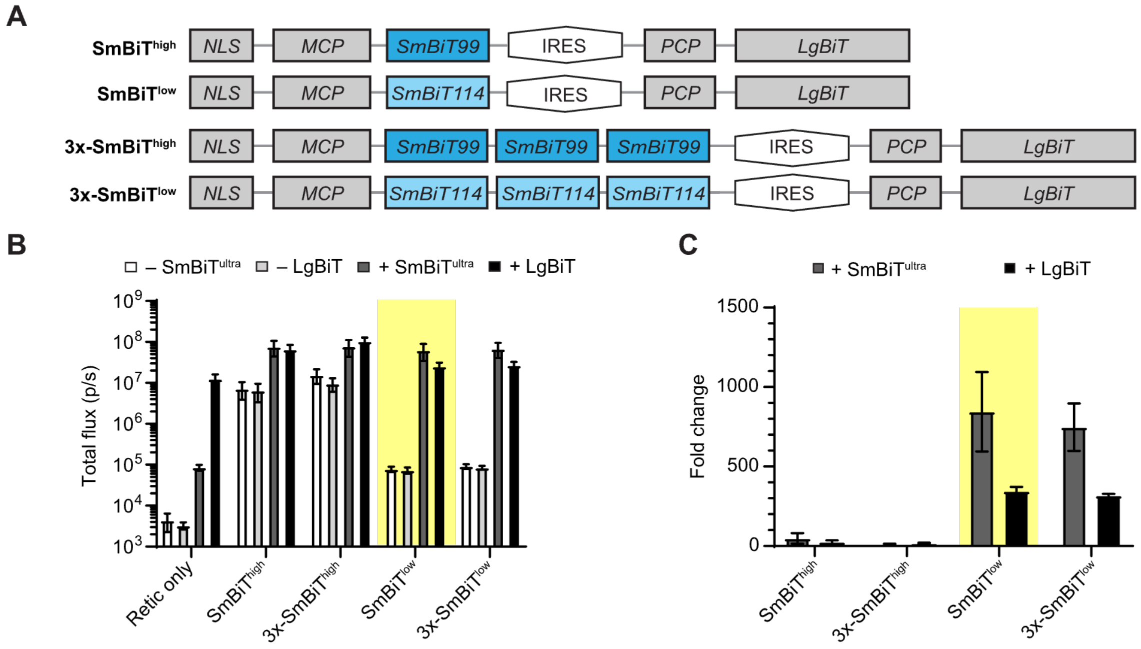


**Optimization of the split Nluc RNA lantern.** (**A**) SmBiT99 (SmBiT$^{high}$, dark blue) and SmBiT114 (SmBiT$^{low}$, light blue) and their serial trimers were constructed as NLS-MCP fusions and co-expressed with IRES-driven PCP-LgBiT. (**B**) The lantern constructs were expressed in IVTT (rabbit reticulocyte lysate, RRL) and evaluated for RNA-independent photon production alone or supplemented with synthetic SmBiT86 (SmBiT$^{ultra}$ ,10 μM) or recombinant LgBiT (10 μM), as noted. The IVTT-expressed lanterns provide background photon measurements, whereas the supplemented experiments (+SmBiT$^{ultra}$ or + LgBiT) represent the saturated complexes, reporting the maximum potential photon output from each construct. (**C**) Fold change in light output from (**B**), calculated for pairs of (saturated/IVTT-expressed) constructs. Error bars represent the standard error of the mean (SEM) for $n$ = 3 replicates experiments.

**Fig. S2.**

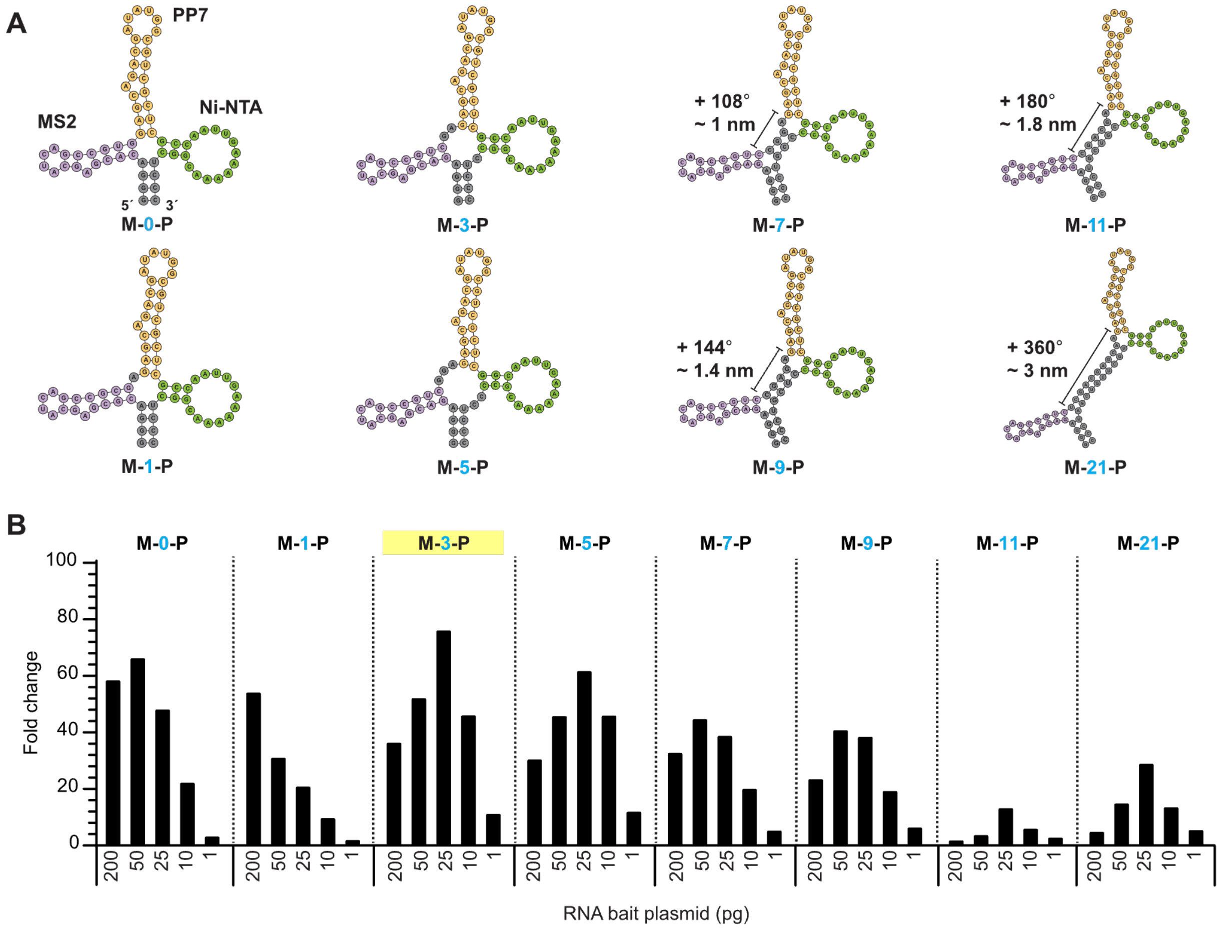


**Design of novel MCP/PCP baits.** (**A**) RNA baits were designed with varying number of nucleotides between the MS2 (purple) and PP7 (orange) aptamers. A Ni-NTA aptamer (green) was also added for purification purposes and improved structural rigidity. (**B**) Luminescence readouts obtained with each RNA bait. Data are plotted as the fold change in signal from each RNA bait over a no-RNA control sample. Decreasing concentrations of RNA bait template were assessed with 1 ng of the lantern DNA plasmid. The mean fold change for each sample ($n$ = 2) is shown.

Fig. S3.

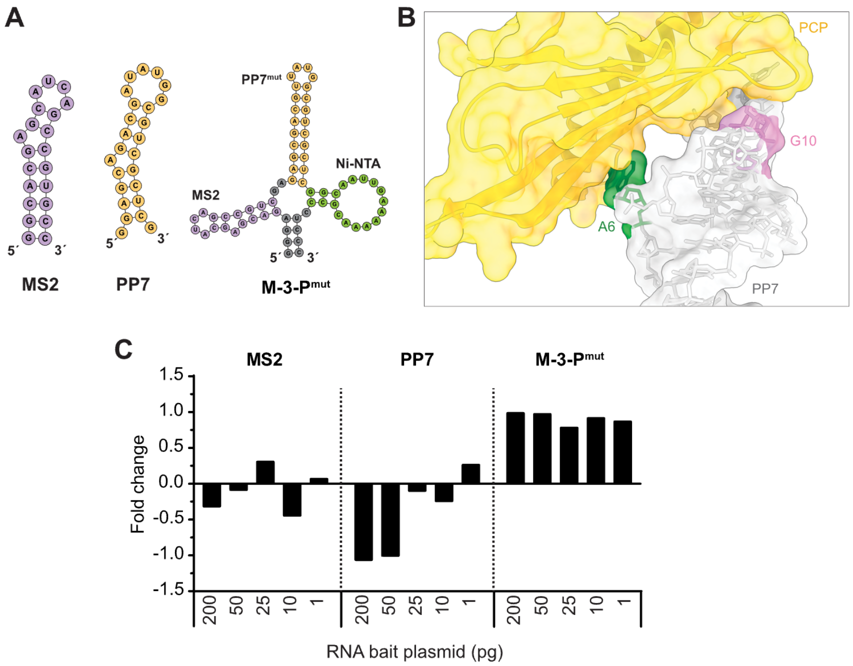


**Lack of luminescence observed with control RNAs.** (**A**) Predicted secondary structures of control RNA baits comprising individual MS2 or PP7 aptamers or the M-3-P$^{mut}$ construct containing two mutations (deletion of A6 and G10). (**B**) Location of the deleterious PP7 mutations in the structure of the unmutated aptamer-protein complex (PDB: 2QUX). (**C**) Fold change in signal observed with each control RNA bait over a no-RNA sample. Decreasing concentrations of RNA bait template were tested with 1 nM of the lantern DNA plasmid. The mean fold change for each sample ($n$ = 2) is shown.

**Fig. S4.**

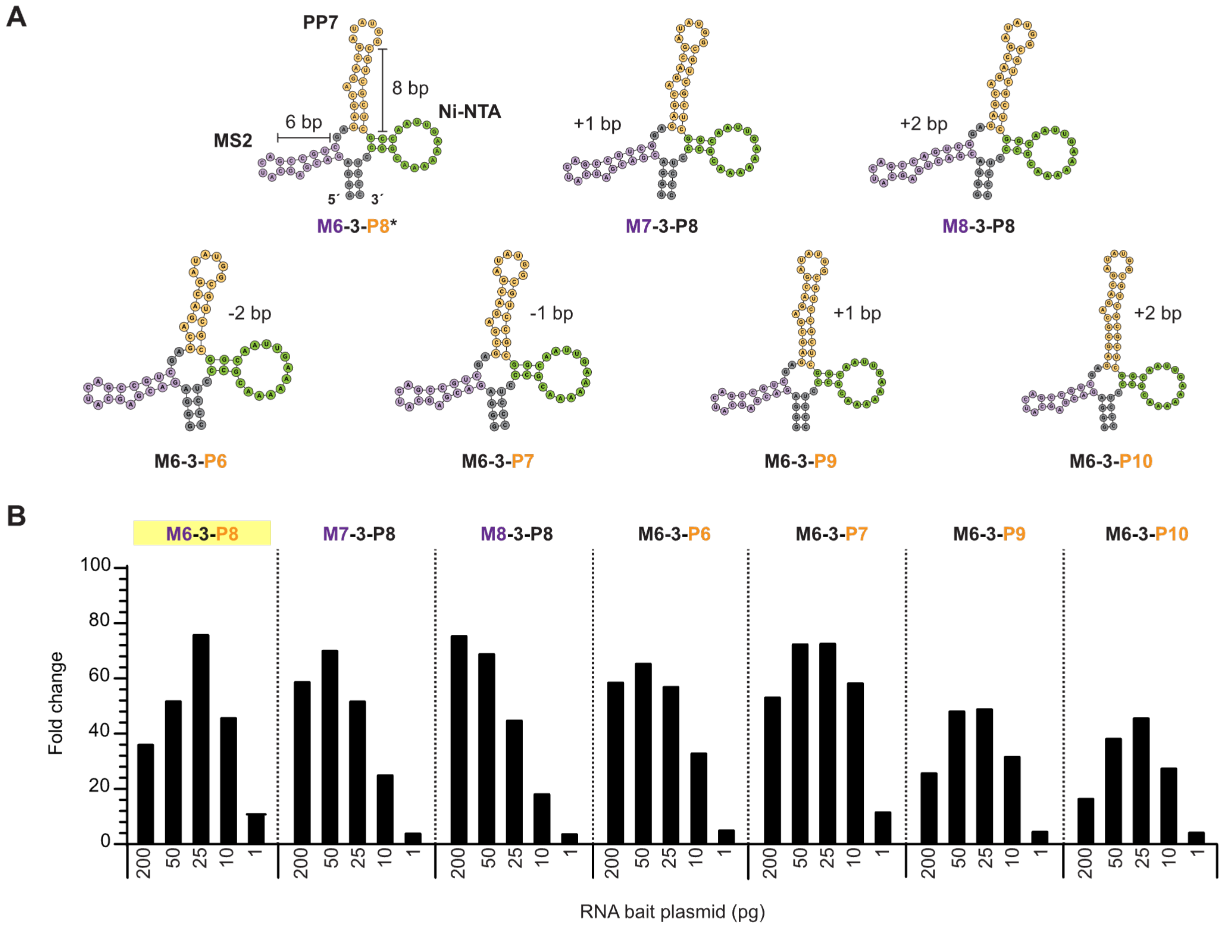


**Modulation of aptamer stem lengths.** (**A**) The stem regions of the MS2 (purple) and PP7 (orange) aptamers were varied by changing the number of base pairs. The asterisk indicates RNA bait with unmodified MS2 and PP7 stems (Fig. 1E). (**B**) Fold change in signal observed with each RNA bait design over a no-RNA control. Decreasing concentrations of each RNA bait template were tested with 1 ng of the lantern DNA plasmid. The mean fold change for each sample ($n$ = 2) is shown.

**Fig. S5.**

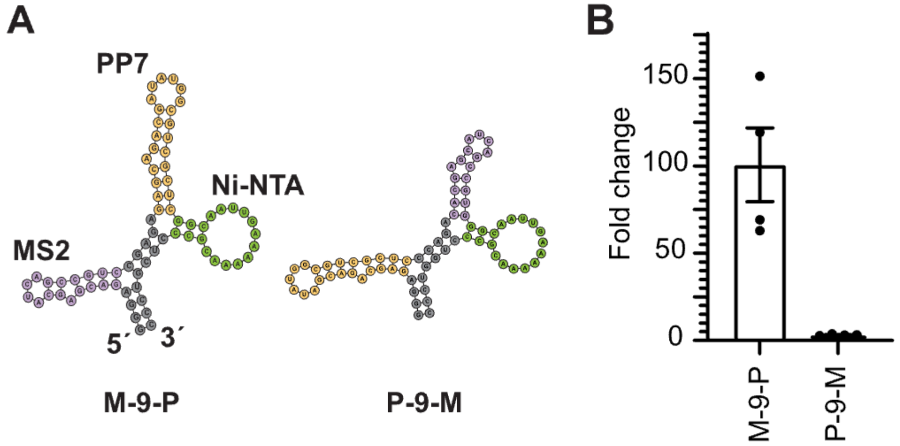


**Aptamer order is critical for lantern assembly**. (**A**) The PP7 aptamer (orange) was placed either 5′ (right) or 3′ (left) of the MS2 aptamer (purple) to evaluate the effect of binding site orientation on lantern function. (**B**) RNA baits were assessed using IVTT and the fold changes in light output over samples without RNA bait are plotted. Error bars represent the standard error of the mean (SEM) for $n$ = 4 replicates.

**Fig. S6.**

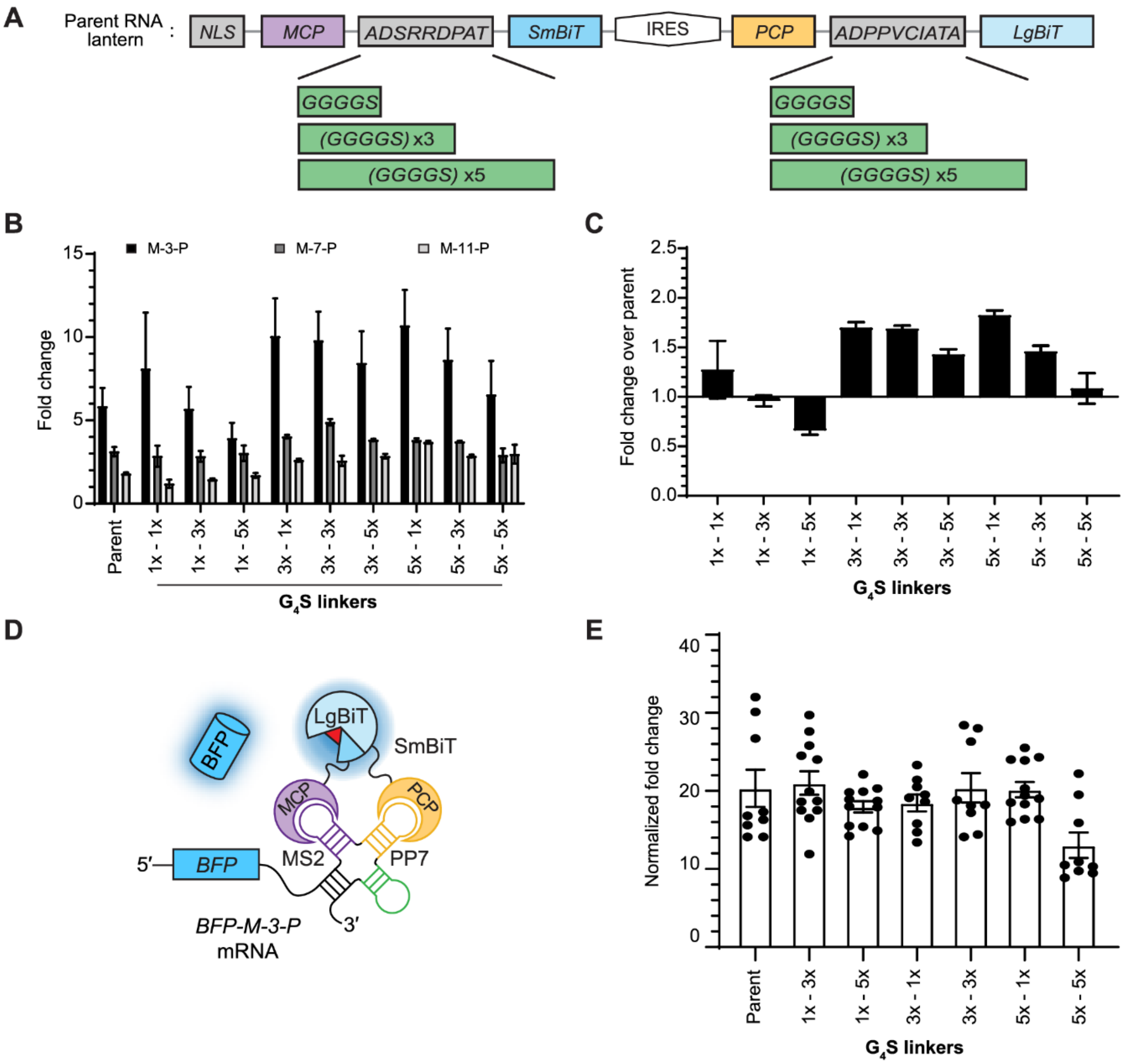


**Lantern linker lengths have a smaller effect on photon output than RNA bait structure.** (**A**) Schematic of G4S linker changes in the RNA lantern. (**B**) Protein linkers comprising 1–5 copies of a glycine-serine repeating unit (G4S) were examined in IVTT. Luminescence outputs were recorded in the presence of various RNA baits. Data are plotted as the fold change in signal over no-RNA bait controls. (**C**) Fold change in luminescence observed with RNA lanterns comprising altered linkers and M-3-P versus the original lantern and M-3-P. No substantial increase in signal was observed with probes comprising G4S units. For (**A**)–(**B**), error bars represent the standard error of the mean (SEM) for $n = 4$ replicates. (**D**) Schematic of an mRNA encoding blue fluorescent protein (BFP) and M-3-P in the 3′ UTR. Transcription of *BFP-M-3-P* mRNA recruits the RNA lantern, resulting in light production. BFP fluorescence enables confirmation of expression. (**E**) RNA lanterns comprising various linker lengths were examined. Cells stably expressing each lantern were transfected with the *BFP-M-3-P* construct. Data are plotted as the fold-change in signal over non-transfected cells. Luminescence measurements were normalized to BFP expression (assessed via flow cytometry). Nine replicates are shown, and the error bars represent the standard error of the mean (SEM) for all replicates. Minimal improvement over the initial RNA lantern design was observed with varying glycine-serine linkers. The 5x-5x design (five $G_4S$ linkers in both protein fusions) exhibited reduced signal turn-on both *in vitro* and *in cellulo*, suggesting disfavored complementation with increasing linker length.

### Fig. S7.

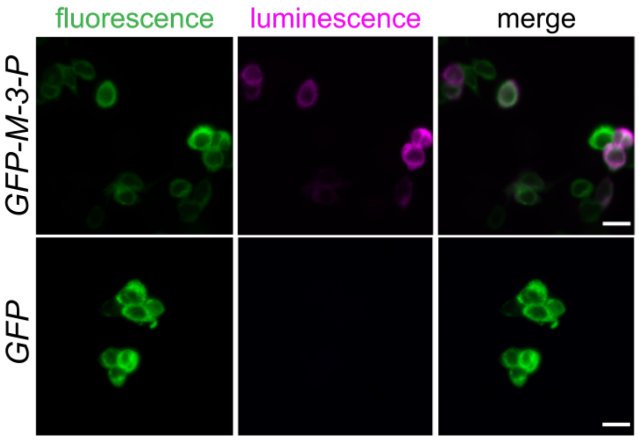


**RNA imaging *in cellulo* with bioluminescent lantern.** HEK293T cells expressing RNA lanterns were transfected with DNA encoding *GFP–M-3-P* (Pearson's r = 0.57) or *GFP* alone (Pearson's r = 0.15). The images shown are replicates of the experiment described in Fig. 4B. Scale bar = 20 µm.

### Table S1. Plasmids used in the study.

| Features | Plasmid Vector | Promoter | Expression in | Used in |
|---|---|---|---|---|
| NLS - HA - MS2 - SmBiT114 - IRES - PP7 - LgBiT - FLAG | pCDNA 3.1/ pLenti 3rd generation | CMV | mammalian | Fig. 1D-G 2B-F Fig. 4B-G S1 S2 S3 S4 S5 S6 S7 |
| NLS - HA - MS2 - SmBiT99 - IRES - PP7 - LgBiT - FLAG | pCDNA 3.1 | CMV | mammalian | S1 |
| NLS - HA - MS2 - 3x SmBiT99 - IRES - PP7 - LgBiT - FLAG | pCDNA 3.1 | CMV | mammalian | S1 |
| SV40 NLS - HA - MS2 - 3x SmBiT114 - IRES - PP7 - LgBiT - FLAG | pCDNA 3.1 | CMV | mammalian | S1 |
| NLS - HA - MS2 - G4S - SmBiT114 - IRES - PP7 - G4S - LgBiT - FLAG | pCDNA 3.1 | CMV | mammalian | S6 |
| NLS - HA - MS2 - G4S - SmBiT114 - IRES - PP7 - G4Sx3 - LgBiT - FLAG | pCDNA 3.1 | CMV | mammalian | S6 |
| NLS - HA - MS2 - G4S - SmBiT114 - IRES - PP7 - G4Sx5 - LgBiT - FLAG | pCDNA 3.1 | CMV | mammalian | S6 |
| NLS - HA - MS2 - G4Sx3 - SmBiT114 - IRES - PP7 - G4S - LgBiT - FLAG | pCDNA 3.1 | CMV | mammalian | S6 |
| NLS - HA - MS2 - G4Sx3 - SmBiT114 - IRES - PP7 - G4Sx3 - LgBiT - FLAG | pCDNA 3.1 | CMV | mammalian | S6 |
| NLS - HA - MS2 - G4Sx3 - SmBiT114 - IRES - PP7 - G4Sx5 - LgBiT - FLAG | pCDNA 3.1 | CMV | mammalian | S6 |
| NLS - HA - MS2 - G4Sx5 - SmBiT114 - IRES - PP7 - G4S - LgBiT - FLAG | pCDNA 3.1 | CMV | mammalian | S6 |
| NLS - HA - MS2 - G4Sx5 - SmBiT114 - IRES - PP7 - G4Sx3 - LgBiT - FLAG | pCDNA 3.1 | CMV | mammalian | S6 |
| NLS - HA - MS2 - G4Sx5 - SmBiT114 - IRES - PP7 - G4Sx5 - LgBiT - FLAG | pCDNA 3.1 | CMV | mammalian | S6 |
| NLS - HA - MS2 - linker - SmBiT114 - IRES - PP7 - linker - YeLgBiT - FLAG | pCDNA 3.1 | CMV | mammalian | Fig. 3G-H |
| NLS - HA - MS2 - linker - SmBiT114 - IRES - PP7 - linker - LumiLgBiT - FLAG | pCDNA 3.1 | CMV | mammalian | Fig. 3G-H |
| NLS - HA - MS2 - FlucN - IRES - PP7 - FlucC - FLAG | pCDNA 3.1 | CMV | mammalian | Fig. 3C |
| BFP - MS2-3-PP7 | pCDNA 3.1 | CMV | mammalian | Fig. 4F-G S6 |
| BFP - MS2-3-PP7 (mut) | pCDNA 3.1 | CMV | mammalian | Fig. 4F-G |
| Staygold - MS2-3-PP7 | pCDNA 3.1 | CMV | mammalian | Fig. 3D-E S7 |
| Staygold | pCDNA 3.1 | CMV | mammalian | Fig. 3D-E S7 |
